# Regional genomic heritability mapping for agronomic traits in sugarcane

**DOI:** 10.1101/2020.04.16.045310

**Authors:** Pedro Marcus Pereira Vidigal, Mehdi Momen, Paulo Mafra de Almeida Costa, Márcio Henrique Pereira Barbosa, Gota Morota, Luiz Alexandre Peternelli

## Abstract

**Background:** The identification of genomic regions involved in agronomic traits is the primary concern for sugarcane breeders. Genome-wide association studies (GWAS) leverage the sequence variations to bridge phenotypes and genotypes. However, their effectiveness is limited in species with high ploidy and large genomes, such as sugarcane. As an alternative, a regional heritability mapping (RHM) method can be used to capture genetic signals that may be missed by GWAS by combining genetic variance from neighboring regions. We used RHM to screen the sugarcane genome aiming to identify regions with higher heritability associated with agronomic traits. We considered percentage of fiber in sugarcane bagasse (FB), apparent percentage of sugarcane sucrose (PC), tonnes of pol per hectare (TPH), and tonnes of stalks per hectare (TSH).

**Methods:** Sequence-capture data of 508 sugarcane (*Saccharum* spp.) clones from a breeding population under selection were processed for variant calling analysis using the sugarcane genome cultivar R570 as a reference. A set of 375,195 single nucleotide polymorphisms were selected after quality control. RHM was conducted by splitting the sugarcane genome into windows of 2 Mb length.

**Results:** We selected the windows explaining > 20% of the total genomic heritability for TPH (64 windows - 5,654 genes) and TSH (72 windows - 6,050 genes), and > 15% for PC (16 windows - 1,517 genes) and FB (17 windows - 1,615 genes). The top five windows that explained the highest genomic heritability ranged from 20.8 to 24.6% for FB (629 genes), 18.0 to 22.0% for PC (452 genes), 53.8 to 66.0% for TPH (705 genes), and 59.5 to 67.4% for TSH (413 genes). The functional annotation of genes included in those top five windows revealed a set of genes that encode enzymes that integrate carbon metabolism, starch and sucrose metabolism, and phenylpropanoid biosynthesis pathways.

**Conclusions:** The selection of windows that explained the large proportions of genomic heritability allowed us to identify genomic regions containing a set of genes that are related to the agronomic traits in sugarcane. These windows spanned a region of 58.38Mb, which corresponds to 14.28% of the reference assembly in the sugarcane genome. We contend that RHM can be used as an alternative method for sugarcane breeders to reduce the complexity of the sugarcane genome.

## 1. Introduction

Sugarcane (*Saccharum* spp.) is a perennial C_4_ grass of the Poaceae family, which is an economically important crop for the sugar and biofuels industries. Its cultivars are multiplied through vegetative propagation and are primarily grown in tropical and subtropical regions (Barbosa et al., 2012).

Improvement of sugarcane is a challenging process guided by conventional methods, which demand time and is hindered by its genomic complexity (Barbosa et al., 2012). Due to its long production cycle, sugarcane breeding programs take at least 11 years to release a new cultivar (Barbosa et al., 2012; Peternelli et al., 2018). Sugarcane breeders also must deal with the complexity of the sugarcane genome that exceeds other crops, which is the product of interspecific hybridization between *Saccharum officinarum* and *S. spontaneum* originated from modern cultivars. These cultivars are highly heterozygous, aneuploid, and have large genomes with a variable number of chromosomes (Lu et al., 1994; D’Hont et al., 2001; Piperidis, Piperidis & D’Hont, 2010; Garsmeur et al., 2018).

High throughput sequencing technologies have been enabling significant advances in uncovering and understanding the genomic complexity of sugarcane. A comparison of the sugarcane genetic maps with other Poaceae species maps revealed microsynteny with the *Sorghum bicolor* genome (Figueira et al., 2012; Aitken et al., 2014; Yang et al., 2017). Analysis of transcriptomes further confirmed that sugarcane and sorghum transcripts share more than 90% of sequence identity (Nishiyama et al., 2014; Yang et al., 2017). Thus, the sorghum genome became the most suitable reference for sequence variation analysis in sugarcane and is of frequent use in linkage and quantitative trait loci (QTL) mapping. However, the recently released reference sequence of the sugarcane cultivar R570 monoploid genome (Garsmeur et al., 2018) opens a new opportunity for sugarcane geneticists. This reference is a high-quality sequence that represents the gene space of the sugarcane monoploid genome, which contains genes annotated in their genomic context with the respective regulatory elements.

Identification of genomic regions involved in agronomic traits related to yield and disease resistance by performing molecular marker-assisted selection is the primary concern for sugarcane breeders. To increase the resolution of single nucleotide polymorphism (SNP) genotyping, sequence-capture methods have been used as a powerful tool to assess sequence variation in target regions across the genomes and to reduce costs in comparison with whole-genome sequencing. Genome-wide association studies (GWAS) leverage these sequence variations to bridge phenotypes and genotypes. GWAS also exploit the high amount of linkage disequilibrium (LD) in sugarcane as a more suitable alternative for bi-parental QTL mapping (Raboin et al., 2008). However, many instances of linked markers still will not be recognized due to the confounding effect of polyploidy of the sugarcane genome (Raboin et al., 2008).

Additionally, GWAS in high ploidy species, such as sugarcane, is also limited by high sequencing depth needed to call variants from the large genome and the difficulty in determining the dosage of markers (Meirmans, Liu & Van Tienderen, 2018). For instance, only a few significantly associated SNPs have been detected in agronomic traits (Yang et al., 2018; Fickett et al., 2019). As an alternative, a regional heritability mapping (RHM) method can be used to capture genetic signals that may be missed by GWAS by combining genetic variance from neighboring regions (Nagamine et al., 2012; Shirali et al., 2016; Resende et al., 2018). Here, we analyzed sequence-capture data of sugarcane clones from a breeding population under selection, which were phenotypically evaluated. We used the RHM method to identify genomic regions that explain significant additive genetic variance in agronomic traits by screening the whole sugarcane genome.

## 2. Materials and Methods

### 2.1. Plant material

The sugarcane clones analyzed in this study were selected from a breeding population of the Sugarcane Breeding Program (Programa de Melhoramento da Cana-de-Açúcar, PMGCA) at the Federal University of Viçosa (Universidade Federal de Viçosa, UFV). This population consisted of 508 clones from 100 half-sib families, originated from crossings between elite sugarcane clones and commercial cultivars.

These crossings were performed in 2010 at the Serra do Ouro’s experimental station located in Murici (Alagoas, Brazil) (09°13’ S, 35°50’ W, 450 m altitude). The seedlings were produced at PMGCA’s experimental station located in Oratórios (Minas Gerais, Brazil) (20°25’ S, 42°48’ W, 494 m altitude). The seedlings were conducted in the first phase trials (T1) performed in 2011 (plant cane) and 2012 (first ratoon). Sugarcane clones analyzed were originated from plants selected in T1 and advanced to the second phase trial (T2) in July 2012, which was conducted in an augmented block design experiment (Federer & Raghavarao, 1975). In the T2 trial, the experimental plots consisted of one 4 m row with a spacing of 1 m between plots. The clones (unreplicated) and two common reference cultivars (checks replicated once each) were arranged in 49 augmented blocks. The reference cultivars were RB867515 (Barbosa et al., 2001) and SP80-1842, which are cultivated in large areas and commonly used as checks in breeding experiments in Brazil.

### 2.2. Phenotypic data

Sugarcane clones were phenotypically evaluated in plant cane in July 2013 after 12 months of growth. Ten stalks randomly taken from each plot were used for estimating the tonnes of stalks per hectare (TSH), percentage of fiber in sugarcane bagasse (FB), apparent percentage of sucrose in sugarcane (PC), and tonnes of pol per hectare (TPH).

TSH was obtained from the total number of stalks per row and the wet weight of 10 stalks determined with a dynamometer (Castro et al., 2016). FB was determined on a wet basis from a 500 g sample of the shredded stalk (Tanimoto, 1964; Legendre, 1992) as [FB(%) = ((100 × DM) – (WM × Brix))/(5 × (100 - Brix))], where DM and WM are dry and wet mass of the sample removed from a hydraulic press and Brix is the juice Brix measured by refractometer. The apparent percentage of sucrose in juice (polarization, POL) was measured by polarimetric determination after juice extraction from 500 g samples crushed in a hydraulic press (Schneider, 1979) and used to derive the apparent percentage of sucrose in sugarcane (PC), according to the following expression (Baffa et al., 2014): [PC (%) = POL × (1 - 0.01 × FB) × (0.9961 - 0.0041 × FB)]. The trait TPH, expressed as a percentage of apparent sucrose on a fresh weight basis, was estimated as [TPH (%) = (TSH × PC) / 100].

### 2.3. Statistical analysis of phenotypic data

We analyzed the phenotypic data through the mixed model methodology using the software Selegen-REML/BLUP (Resende, 2016). Variance components were estimated by restricted maximum likelihood (REML), and the genotypic effects of the clones were predicted by BLUP. We used the following linear mixed model:

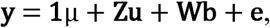

where **y** is the vector of phenotypic observations for each trait; **1** is a vector of 1s; µ is the overall mean; **u** ∼ N(**0, I**σ^2^_*u*_ ) is the vector of random genotypic effects; **b** ∼ N(**0, I**σ^2^_*b*_) is the vector of random block effects; and **e** ∼ N(**0, I**σ _*e*_) is the vector of residuals. **Z** and **W** are incidence matrices relating the observations to the respective model effects.

The statistical significance of the effects was tested using the likelihood ratio test under the analysis of deviance theory (Resende, 2016). The predicted genotypic values 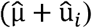 of the clones (*i* = 1 to n) are listed in Table S1 and were considered as response variables in the RHM analysis.

### 2.4. Genotypic data: Genomic DNA sequencing

The total genomic DNA of sugarcane clones was extracted using the DNeasy Plant Mini Kit of Qiagen^®^ following the manufacturer’s guidelines. The genomic libraries were produced and sequenced by RAPiD Genomics (Florida, USA). In this sequencing, the single-end libraries were built using a capture-seq methodology (Neves et al., 2013), which includes a set of probes to capture non-repetitive and evenly distributed sequences in the sugarcane genome.

Briefly, a set of 50,000 unique sequences was identified from: i) existing expressed sequence tags (ESTs) from public sugarcane cDNA libraries and ii) whole-shotgun genome sequences available publicly, consistently distributed in the genome and assuming synteny to the sorghum genome. Biotinylated 120-mer probes that complement a segment of each of the 50,000 target regions of the sugarcane genome were synthesized and were utilized to capture sequences at each target locus. The sequencing yielded a dataset of 4.77 billion reads containing sequences with 100nt or 150nt in length.

### 2.5. Bioinformatic and genetic analysis of genotypic data

#### 2.5.1 Reference genome

To evaluate the sequence variation, we selected the genome of sugarcane cultivar R570 (Grasmeur et al. 2018) as a reference, which is available at Sugarcane Genome Hub (http://sugarcane-genome.cirad.fr/organism/R570-Sugarcane/cultivar). This genome is a Single Tilling Path (STP) assembly of 408.94Mb containing ten chromosomes and 24,341 annotated genes. The unplaced contigs were not considered for the mapping analysis.

#### 2.5.2 Mapping analysis

The raw reads of each capture-seq library were first trimmed to remove sequence adapters and poorly sequenced regions using Trimmomatic version 0.38 (Bolger, Lohse & Usadel, 2014). Trimmed reads were mapped to the reference genome using the BWA-MEM algorithm of BWA version 0.7.17 (http://bio-bwa.sourceforge.net/) (Li & Durbin, 2009). A flag identifying the several sugarcane clones was added to each mapping file. Then, the Sequence Alignment Map files were processed using SortSam, MarkDuplicates, and BuildBamIndex tools in Picard version 2.18.27 (https://github.com/broadinstitute/picard/). As a result, we produced ordered and deduplicated Binary Alignment Map files, containing ordered and deduplicated data. A schematic overview of the analysis conducted in the current study is shown in Figure S1.

#### 2.5.3. Variant calling

Variants were called using FreeBayes version 1.2.0 (https://github.com/ekg/freebayes) (Garrison & Marth, 2012) with a minimum mapping quality of 20, minimum base quality of 20, and minimum coverage of 20 reads at every position in the reference genome. After variant calling, SNPs were filtered using vcftools version 0.16.15 (https://vcftools.github.io/index.html), Bcftools version 1.9 (https://samtools.github.io/bcftools/), and in-house AWK shell scripts. Among the polymorphic loci detected, we selected those with biallelic SNPs with less than 25% of missing data. Missing genotypes were imputed by Beagle version 5.1 (Browning, Zhou & Browning, 2018), using a flexible localized haplotype-cluster model to group locally similar haplotypes into clusters.

#### 2.5.4. Regional heritability mapping (RHM) analysis

We performed RHM based on the variance component method described in Nagamine et al. (2012) using REACTA (Regional Heritability Advanced Complex Trait Analysis) version 0.97 (Canela-Xandri et al., 2015). We concatenated the ten chromosomes and split the whole sugarcane genome into 409 overlapping windows with an average length of 2 Mb to estimate the proportion of phenotypic variance explained by all genome-wide SNPs or a subset of SNPs using REML. The following mixed model is considered:

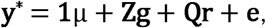

where **y**^*^ is the vector of adjusted phenotypic observations (genotypic values), **1** is a vector of 1s, µ is the overall mean, **Z** and **Q** are the design matrices for the whole (without the window) and the regional random effects, respectively. The distributions and covariance structures of **g** and **r** were **g** ∼ N(**0, G**σ^2^_*g*_) and **r** ∼ N(**0, G**_*r*_σ^2^_*r*_), respectively. The residual term followed **e** ∼ N(**0, I**σ _*e*_). We also run a model using all genome-wide SNPs without any window to estimate the total genomic heritability for each trait (h^2^_*G*_).

A Bonferroni correction based on the number of independent windows was used to obtain a genome-wide significance threshold for the RHM analysis, [αcritical = 0.05/(0.5 x Num. Windows)], as previously proposed (Nagamine et al., 2012) and implemented (Shirali et al., 2016) in the literature.

#### 2.5.5. Functional analysis of genomic windows

Among the analyzed genomic regions, we selected the windows explaining above 20% of the total genomic heritability for TPH and TSH (h^2^_*r*_ */* h^2^*G*≥ 20%) and above 15% for PC and FB (h^2^_*r*_ */* h^2^*G* ≥ 15%), where h^2^*r* = σ^2^_*r*_ / (σ^2^_*g*_ + σ^2^_*r*_ + σ^2^_*e*_) and h^2^_*G*_ = σ^2^*g* / (σ^2^_*g*_+ σ^2^_*r*_+ σ^2^_*e*_). The gene content of these windows was identified using bedtools version 2.28.0 and the gff3 genome annotation file of the sugarcane cultivar R570 genome. The protein sequences encoded by the genes included in those windows were functionally characterized through similarity searches using BLAST version 2.6.0 (Altschul et al., 1990), Blast2GO (Gotz et al., 2008), and KAAS (KEGG Automatic Annotation Server) (Moriya et al., 2007). The lists of genes that explained high heritability for each trait were compared through Venn diagrams using the jvenn package (Bardou et al., 2014). Additionally, the gene ontology (GO) terms of these genes were further summarized using REViGO (Supek et al., 2011).

## 3. Results

### 3.1. Phenotypic variation of analyzed traits

Phenotypic analysis of the sugarcane clones showed a considerable phenotypic variation for the traits analyzed suggesting that there is genotypic variability to be exploited (Figure 1; Tables S1 and S2). The genotypic values obtained with the adjusted model, described above, ranged from 8.39 to 16.43 (average of 11.09) for a percentage of FB, from 6.50 to 17.00 (av. 13.81) for PC, from 6.81 to 20.66 (av. 14.83) for TPH, and from 51.52 to 152.23 (av. 107.96) for TSH (Table S1). Genotypic variance (σ^2^_u_) and broad-sense heritability (H^2^) were 2.21 and 0.56 for FB; 2.02 and 0.71 for PC; 7.48 and 0.58 for TPH; 389.98 and 0.57 for TSH, respectively (Table S2).

**Figure 1.**
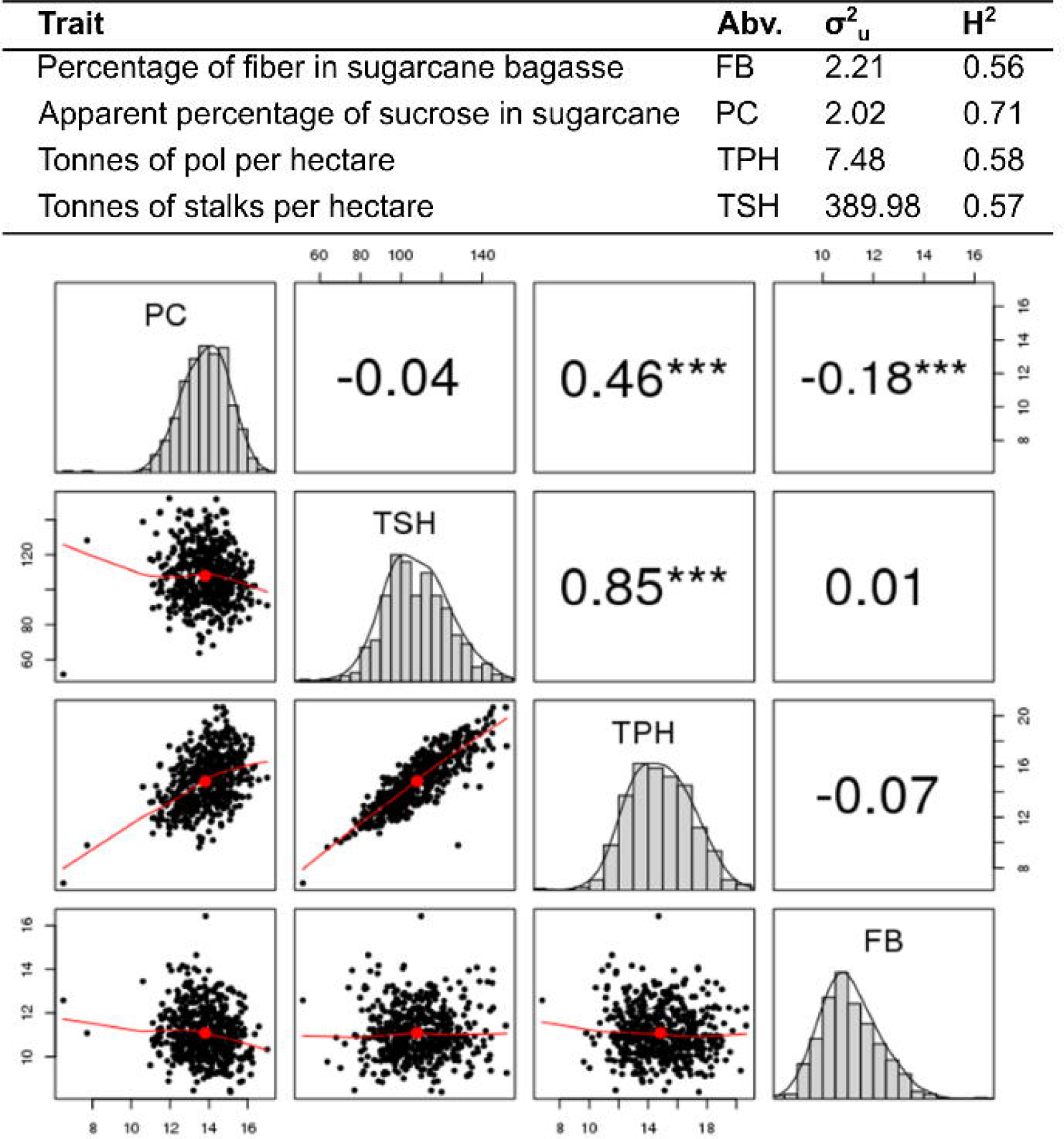
Overview of genotypic values (overall mean + BLUP) of traits evaluated in the sugarcane breeding population. Five hundred eight sugarcane clones were evaluated for percentage of fiber in sugarcane bagasse (FB), apparent percentage of sucrose in sugarcane (PC), tonnes of pol per hectare (TPH), and tonnes of stalks per hectare (TSH). σ^2^_u_: genotypic variance. H^2^: broad-sense heritability. ***: P-value < 0.01.

Pearson’s pairwise correlations between adjusted genetic values of FB, PC, TPH, and TSH are shown in Figure 1 and evaluated for their significance (P-value ≤ 0.01). We observed FB and PC were negatively correlated (−0.18). TPH was positively correlated with PC (0.46) and highly correlated with TSH (0.86). FB was not significantly correlated with TSH and TPH. We observed a weak and negligible correlation between TSH and PC.

### 3.2. Genotyping analysis and variant selection

Among all 4.77 billion reads sequenced from the 508 sugarcane clones, 4.58 billion (96.01%) were trimmed and selected for being mapped on the reference genome (∼ 9.02 million of reads per sugarcane clone). These reads were mapped with an average rate of 75.73% (ranging from 71.39 to 77.94% among the sugarcane clones) and genome coverage of 1.46X (ranging from 1.14 to 1.88 among the sugarcane chromosomes) (Table 1).

**Table 1.** Summary of genotyping of analyzed sugarcane genotypes. Capture sequencing data of 508 sugarcane genotypes were mapped to the reference genome, and variant calling analysis was performed with the SNP quality control process.

| Sugarcane chromosome | Length (Mb) | Number of Genes | Average of mapping coverage | Number of SNPs before QC | Number of SNPs after QC |
| --- | --- | --- | --- | --- | --- |
| Sh01 | 72.29 | 4,476 | 1.88 | 127,817 | 59,783 |
| Sh02 | 52.64 | 3,178 | 1.55 | 95,999 | 44,429 |
| Sh03 | 54.89 | 3,399 | 1.46 | 115,485 | 52,545 |
| Sh04 | 45.58 | 2,791 | 1.54 | 101,102 | 46,756 |
| Sh05 | 23.28 | 1,233 | 1.14 | 39,330 | 17,993 |
| Sh06 | 35.41 | 2,108 | 1.79 | 74,477 | 34,853 |
| Sh07 | 30.10 | 1,824 | 1.38 | 63,170 | 27,829 |
| Sh08 | 24.32 | 1,307 | 1.25 | 49,150 | 23,001 |
| Sh09 | 36.58 | 2,082 | 1.32 | 76,803 | 35,419 |
| Sh10 | 34.65 | 1,943 | 1.33 | 71,654 | 32,587 |
| **Total** | 408.94 | 24,341 | 1.46 | 814,987 | 375,195 |

From those reads, 15.41 million polymorphic loci were identified among the sugarcane clones, and 814,987 SNPs were subsequently filtered by selecting biallelic loci with less than 25% of missing data. After SNP imputation and quality control, 375,195 SNPs were selected for the RHM analysis, which corresponds to a frequency of 917.48 SNPs/Mb in the sugarcane genome (Table 1). Among sugarcane chromosomes (Sh), Sh05 was the least polymorphic (772.81 SNPs/Mb), and Sh04 was the most polymorphic (1,025.88 SNPs/Mb).

### 3.3. Regional heritability mapping analysis

The calculated total genomic heritability (h^2^_*G*_) was 0.798 ± 0.233 for FB, 0.932±0.133 for PC, 0.369 ± 0.256 for TPH, and 0.383 ± 0.210 for TSH. A suggestive significance threshold of 3.61 for -log_10_ P-value (i.e., P-value = 0.000244) was calculated for window selection in the RHM analysis. None of the analyzed windows were associated with the traits with P-values above this threshold (Figure S2). Therefore, the selection of windows was proceeded based on window heritability (h^2^_*r*_), and the proportion of h^2^_*G*_ explained (h^2^_*r*_ */* h^2^_*G*_) aiming to identify regions that could be further exploited for prospecting genes (Figure 2).

**Figure 2.**
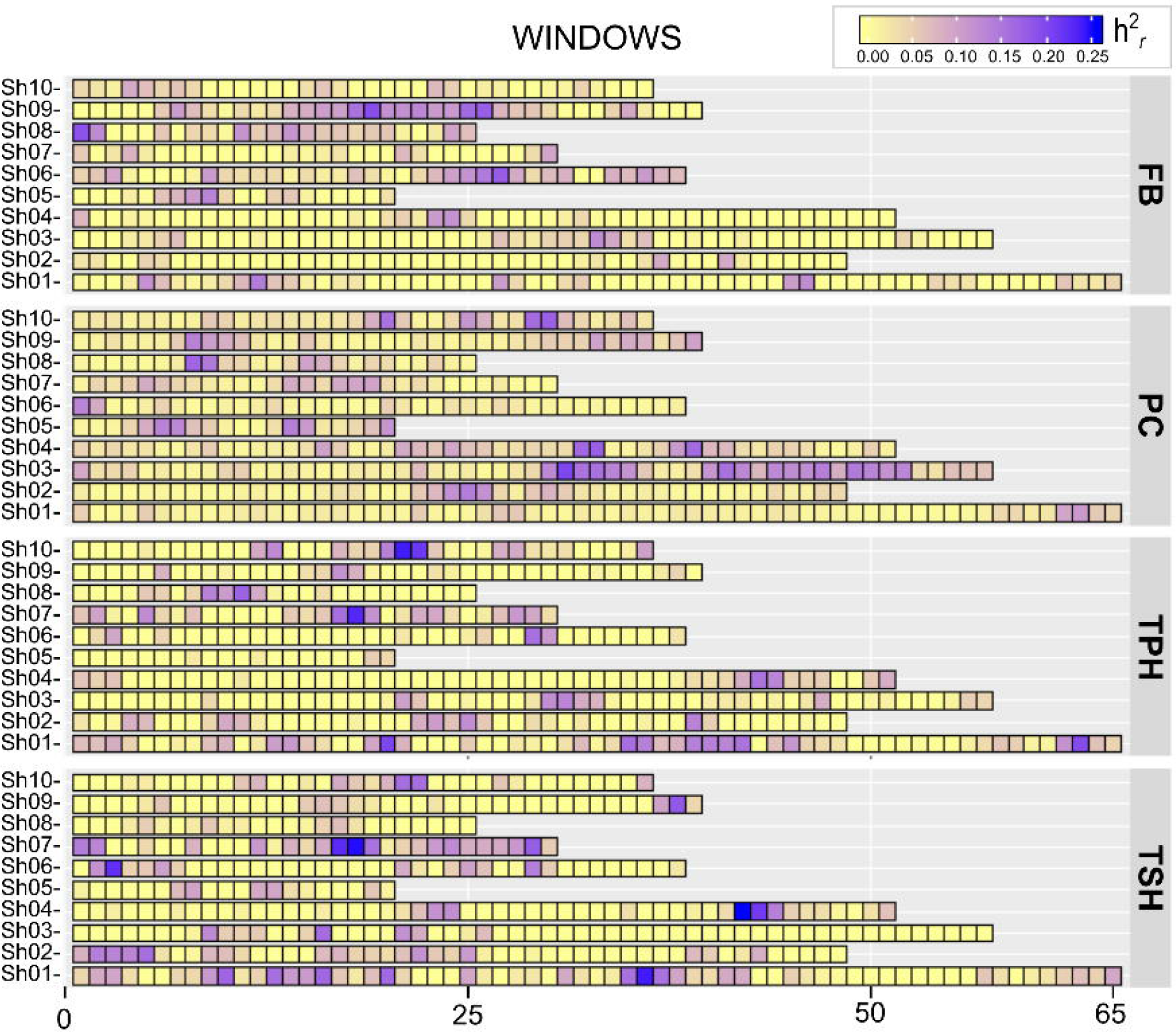
Regional heritability mapping (RHM) analysis of the sugarcane genome. The tilemap of RHM distribution along the genome of 508 sugarcane clones highlights the genomic regions with higher heritability for the percentage of fiber in sugarcane bagasse (FB), apparent percentage of sucrose in sugarcane (PC), tonnes of pol per hectare (TPH), and tonnes of stalks per hectare (TSH). The ten chromosomes sugarcane genome (Sh01 to Sh10) were concatenated and split into windows with 2 Mb length. The color scale on the top-right side of the plot shows the magnitude of h^2^_*r*_ for each window.

Among the analyzed windows, we selected those that explained 15% of h^2^_*G*_ for FB (17 windows - 1,615 genes) and PC (16 windows - 1,517 genes), and those that explained 20% of h^2^_*G*_ for TPH (64 windows - 5,654 genes) and TSH (72 windows - 6,050 genes) (Figures 2 and 3; Table S3). Even though distributed across all the chromosomes, none of these selected windows was related to all traits jointly. One window was related to FB, PC, and TPH (w146 Sh03: 31,787,519 to 34,081,415 - 137 genes), and two windows were related to PC, TPH, and TSH (w90 - Sh02: 25,073,847 to 27,836,987 - 153 genes; w393 - Sh10: 19,693,048 to 22,127,732 - 85 genes). TPH and TSH were the traits that shared the highest number of windows (32 windows - 3,043 genes).

**Figure 3.**
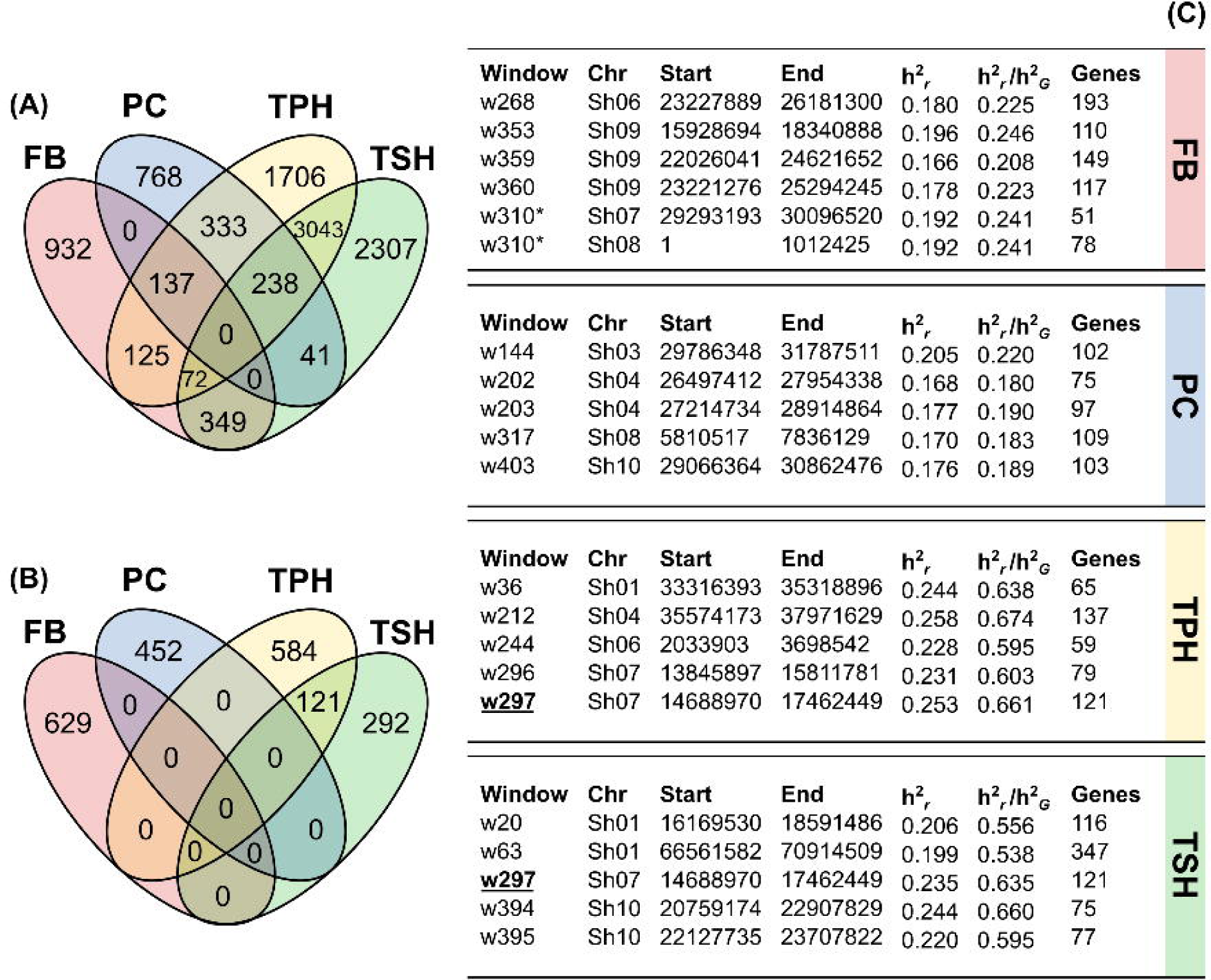
Gene content of sugarcane genomic regions with higher heritability for FB, PC, TSH, and TPH traits in regional heritability mapping analysis. **A)** Gene counting of all regions with higher heritabilities. All genes are listed in Table S3. **B)** Gene counting of top-5 windows with higher heritability. All genes are listed in Table S4. **C)** Detailed information about the top-5 windows. Window 297 is the only one shared by two traits and is marked in bold and underlined. Window 310 spans two chromosomes, comprising a region of 2 Mb, which begins in the last window of Sh07 and ends in the first window Sh08, and is marked with an asterisk.

The top five windows with high heritability explained 20.8 to 24.6% of h^2^_*G*_ for FB (629 genes), 18.0 to 22.0% for PC (452 genes), 53.8 to 66.0% for TPH (705 genes), and 59.5 to 67.4% for TSH (413) (Figure 3). Among these windows, w297 (Sh07: 14688970 to 17462449 121 genes) was the only one related to two traits, and it explained 63.5% of the total heritability for TPH and 66.1% for TSH.

Functional annotations of all the 2,078 genes included in those top five windows generated a non-redundant list of the biological process containing 68 GOs for FB, PC and TPH, and 48 GOs for TSH (Figure 4). Also, 712 genes (34.26%) were classified into 244 KEGG ortholog groups, which were mapped to 104 KEGG pathways, including carbon metabolism, starch, and sucrose metabolism, and phenylpropanoid biosynthesis pathways (Table 2).

**Table 2.** Genes located at the top-5 windows with higher heritability and that are related to carbon metabolism, starch, and sucrose metabolism, and phenylpropanoid biosynthesis pathways. Genes that were assigned to the KEGG ortholog groups (KOs) and mapped to the KEGG pathways of carbon metabolism (CM), glycolysis and gluconeogenesis (GG), sucrose, and starch metabolism (SSM), phenylpropanoid biosynthesis (PB), and oxidative phosphorylation (OP).

| Trait | Gene(window) | KO | Enzyme [EC number] | Pathway |
| --- | --- | --- | --- | --- |
| FB | Sh09_g008010(w353) | K00029 | malate dehydrogenase [1.1.1.40] | CM |
| FB | Sh06_g013660(w268) | K00844 | hexokinase [2.7.1.1] | CM, SSM |
| FB | Sh06_g013010(w268) | K13065 | shikimate O-hydroxycinnamoyltransferase [2.3.1.133] | PB |
| FB | Sh09_g011980(w359/w360) | K23260 | scopoletin glucosyltransferase [2.4.1.128] | PB |
| PC | Sh08_g004080(w317) | K01623 | fructose-bisphosphate aldolase [4.1.2.13] | CM |
| PC | Sh04_g016390(w203) | K01834 | phosphoglycerate mutase [5.4.2.11] | CM |
| PC | Sh04_g015250(w202),<br>Sh04_g015260(w202) | K01900 | succinyl-CoA synthetase [6.2.1.4 6.2.1.5] | CM |
| PC | Sh08_g004580(w317) | K02437 | glycine cleavage system H protein | CM |
| PC | Sh06_g014550(w268),<br>Sh06_g014560(w268) | K01792 | glucose-6-phosphate 1-epimerase [5.1.3.15] | GG |
| PC | Sh03_g016860(w144) | K02150 | V-type H<sup>+</sup>-transporting ATPase subunit E | OP |
| PC | Sh08_g003930(w317) | K02155 | V-type H<sup>+</sup>-transporting ATPase 16kDa proteolipid subunit | OP |
| PC | Sh04_g016300(w203) | K01087 | trehalose 6-phosphate phosphatase [3.1.3.12] | SSM |
| PC | Sh10_g015790(w403),<br>Sh10_g015810(w403) | K01187 | alpha-glucosidase [3.2.1.20] | SSM |
| PC | Sh04_g015390(w202),<br>Sh04_g015400 (w202),<br>Sh04_g015410 (w202) | K10775 | phenylalanine ammonia-lyase [4.3.1.24] | PB |
| PC | Sh04_g015380(w202) | K13064 | phenylalanine/tyrosine ammonia-lyase [4.3.1.25] | PB |
| TPH | Sh01_g042660(w063),<br>Sh01_g042670(w063) | K00640 | serine O-acetyltransferase [2.3.1.30] | CM |
| TPH | Sh01_g011250(w020) | K00873 | pyruvate kinase [2.7.1.40] | CM |
| TPH | Sh01_g042590(w063) | K01681 | aconitate hydratase [4.2.1.3] | CM |
| TPH | Sh10_g011930(w394/w395) | K01738 | cysteine synthase [2.5.1.47] | CM |
| TPH | Sh01_g042950(w063) | K03781 | catalase [1.11.1.6] | CM |
| TPH | Sh10_g012040(w394/w395) | K02147 | V-type H<sup>+</sup>-transporting ATPase subunit B | OP |
| TPH | Sh01_g042110(w063) | K01177 | beta-amylase [3.2.1.2] | SSM |
| TPH | Sh01_g041030(w063) | K01904 | 4-coumarate--CoA ligase [6.2.1.12] | PB |
| TPH | Sh10_g012300(w395) | K12355 | coniferyl-aldehyde dehydrogenase [1.2.1.68] | PB |
| TSH | Sh01_g020340(w036) | K00814 | alanine transaminase [2.6.1.2] | CM |
| TSH | Sh01_g020380(w036) | K01601 | ribulose-bisphosphate carboxylase large chain [4.1.1.39] | CM |
| TSH | Sh01_g020520(w036) | K00001 | alcohol dehydrogenase [1.1.1.1] | GG |
| TSH | Sh01_g020600(w036) | K01785 | aldose 1-epimerase [5.1.3.3] | GG |
| TSH | Sh04_g021630(w212) | K01087 | trehalose 6-phosphate phosphatase [3.1.3.12] | SSM |

**Figure 4.**
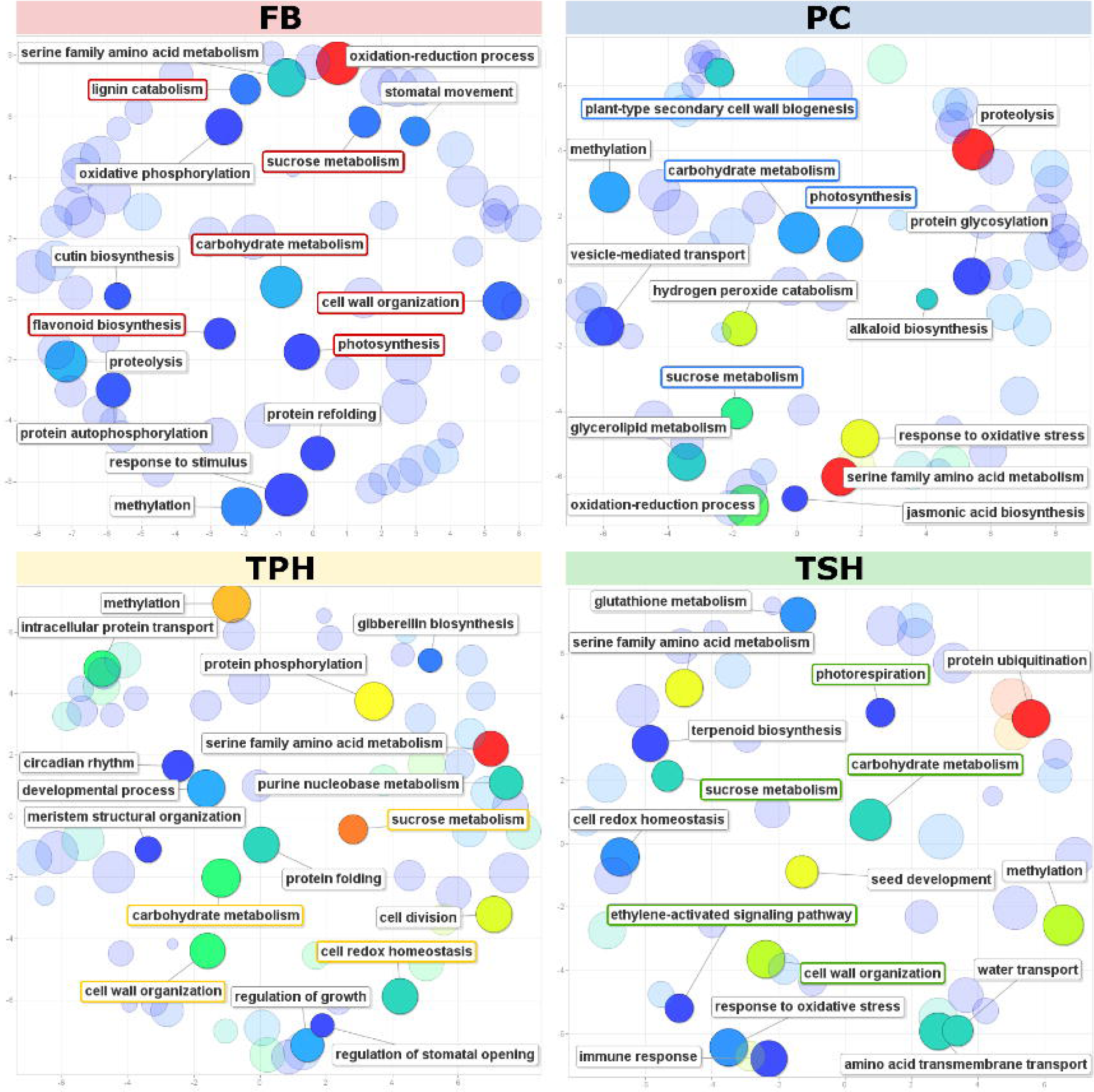
Functional analysis of the genes identified in of top-5 windows of sugarcane genome with higher heritability for FB, PC, TSH, and TPH traits in regional heritability mapping analysis.

## 4. Discussion

Genotyping by sequencing (GBS) coupled with capture-seq methods have been a practical approach to survey polymorphisms in the genomes of *Saccharum* species. It reduces the complexity of genomes and allows genome-wide analysis aiming to develop molecular markers (Song et al., 2016; Balsalobre et al., 2017; Yang et al., 2017, 2018, 2019; Fickett et al., 2019). However, the complexity of the sugarcane genome still poses some challenges for the widespread use of GBS and the adoption of molecular marker-assisted selection in breeding programs. The high and variable ploidy of the allopolyploid genome (Thirugnanasambandam, Hoang & Henry, 2018) makes the identification of SNPs significantly associated with agronomic traits of interest in sugarcane crop a challenging task. The identification of significant SNPs would demand a high coverage when using the GBS approach, which increases the costs involved in population studies (Meirmans, Liu & Van Tienderen, 2018; Yang et al., 2018). Studies reporting SNPs associated with agronomic traits in the sugarcane genome are still scarce. For instance, only 229 SNPs (that explained > 5% phenotypic variation at the unadjusted P-value cutoff of 0.05) have been reported for 97 clones in a breeding population (Fickett et al., 2019) and 191 SNPs (Bonferroni corrected P-value cutoff of 0.05) for 308 accessions in a germplasm collection (Yang et al., 2018).

Based on the low number of SNPs detected at 5% of significance threshold in previous work, and also because we used a low coverage capture-seq data from our breeding population (containing 508 sugarcane genotypes), we performed RHM analysis to investigate regions in the sugarcane genome related to agronomic traits. These sugarcane clones showed comparable phenotypic data, as observed in other studies (Racedo et al., 2016; Yang et al., 2018; Fickett et al., 2019). Their average values of FB, PC, TPH, and TSH were 11.09, 13.81, 14.83, and 107.96, respectively. Comparable TPH of 17.12 t/ha was observed in Reunion Island (Gouy et al., 2014), 9.22 t/ha in Tucumán (Argentina) (Racedo et al., 2016), and 8.84 t/ha in Louisiana (United States) (Fickett et al., 2019).

The high and positive pairwise correlation observed for TPH with TSH and PC is expected because TPH is estimated as a product of these two variables. On the other hand, the negative pairwise correlation between FB and PC is in agreement with the knowledge about carbon partition and metabolism in sugarcane (Hoang et al., 2017). The estimates of broad-sense heritability ranged from 0.56 to 0.71, similar to the range observed for the same traits in other studies with sugarcane (Gouy et al., 2014; Racedo et al., 2016; Fickett et al., 2019).

The RHM analysis appears to be an interesting approach to screening the sugarcane genome aiming to identify regions related to the agronomic traits. We argued that the regions explaining a higher portion of the genomic heritability could be further explored for molecular marker investigation. Unfortunately, in the present study, none of the analyzed genome windows were significantly associated with the traits considered at the suggestive significance threshold of 3.94 (Figure S2), which could be due to the lower sequencing coverage, heterogeneity of regions covered by the sequencing, and the polyploidy effect. To overcome these limitations, we considered a new strategy to infer about the regions of the genome related to the traits under study: the selection of windows showing high heritability (h^2^_*r*_) and proceeded with the analysis of their genic content to identify candidate regions to develop molecular markers. The analyzed traits showed regions with higher heritability, explaining 15% or 20% of h^2^_*G*_ in almost all sugarcane chromosomes. The number of windows above these thresholds was lower for FB (17 windows > 15%; 1,615 genes) and PC (16 windows > 15%; 1,517 genes) when compared to TPH (65 windows > 20%; 5,654 genes) and TSH (73 windows > 20%; 6,050 genes) (Figures 2 and 3). These differences could be due to the lower complexity of FB and PC, which might be controlled by a smaller number of genes. The selection of top-5 windows with higher heritability was enough to reduce the complexity of the sugarcane genome and to provide insights about regions containing a set of genes possibly related to FB (629 genes), PC (452 genes), TPH (705 genes) and TSH (413 genes) traits.

Analysis of biological processes attributed to a function of genes located in the windows with higher heritability for FB (629 genes) indicated GO terms such as “cell wall organization” (GO: 0071555), “flavonoid biosynthetic process” (GO: 0009813), and “lignin catabolic process” (GO: 0046274) (Figure 4). Some of these genes encode enzymes that catalyze the biosynthesis of secondary metabolites such as flavonoids, phenylpropanoids, and lignin. Among these enzymes are shikimate O-hydroxycinnamoyltransferase (window w268; gene Sh06_g013010), scopoletin glucosyltransferase (w359/w360; Sh09_g011980), and flavonoid 3’-monooxygenase (w360; Sh09_g012740). Genes related to “carbohydrate metabolic process” (GO:0005975), “carbon utilization” (GO:0015976), and “sucrose metabolic process” (GO:0005985) are also included in genomic regions with higher heritability for FB. These genes encode enzymes such as hexokinase (w268; Sh06_g013660) and malate dehydrogenase (w353; Sh09_g008010).

Genes related to “carbohydrate metabolic process” (GO:0005975) and “photosynthesis” (GO:0015979) are among those included in the windows with higher heritability for PC (452 genes). These genes encode enzymes of glycolysis/gluconeogenesis, pentose phosphate, oxidative phosphorylation pathways, such as fructose-bisphosphate aldolase (w317; Sh08_g004080), glucose-6-phosphate 1-epimerase (2 copies in w268), phosphoglycerate mutase (w203; Sh04_g016390), succinyl-CoA synthetase (2 copies in w202), V-type proton ATPase subunit E (w144; Sh03_g016860) and vacuolar ATP synthase 16kDa proteolipid subunit (w317; Sh08_g003930). Genes involved in phenylpropanoid biosynthesis pathways, such as phenylalanine ammonia-lyase (PAL) (3 copies in w202), and phenylalanine/tyrosine ammonia-lyase (PTAL) (w202; Sh04_g015380) are also included in these windows.

Windows with higher heritability for TPH (705 genes) contain genes which are related to “carbohydrate metabolic process”, “sucrose metabolic process” (GO:0005985), “cell wall organization” (GO:0071555) and “cell redox homeostasis” (GO:0045454). Pyruvate kinase (w020; Sh01_g011250), aconitate hydratase (w063; Sh01_g042590), vacuolar ATPase B subunit (w394/w395; Sh10_g012040) and beta-amylase (w063; Sh01_g042110) are among the enzymes encoded by these genes, which are part of the carbon, starch, and sucrose metabolism pathways. Also, two genes that encode the enzymes 4-coumarate-CoA ligase (w063; Sh01_g041030) and coniferyl-aldehyde dehydrogenase (w395; Sh10_g012300), which are part of phenylpropanoid biosynthesis pathways, are also included in these high heritability windows.

Among the genes located in windows with higher heritability for TSH (413 genes), some are related to “carbohydrate metabolic process” (GO:0005975), “cell wall organization” (GO:0071555), “sucrose metabolic process” (GO:0005985), “photorespiration” (GO:0009853), and “ethylene-activated signaling pathway” (GO:0009873). The genes encode the enzymes ribulose-1,5-bisphosphate carboxylase/oxygenase large subunit (w036; Sh01_g020380), alanine transaminase (w036; Sh01_g020340), trehalose 6-phosphate phosphatase (w212; Sh04_g021630), alcohol dehydrogenase (w036; Sh01_g020520), aldose 1-epimerase (w036; Sh01_g020600), and ethylene receptor (EIN4) (w244; Sh06_g001310). In contrast to the other traits, none of the windows selected for TSH contain genes related to phenylpropanoid pathways.

Taken together, 14 windows among those top-higher heritabilities for each trait analyzed contain 35 genes that encode enzymes that catalyze biochemical reactions of carbon metabolism, starch and sucrose metabolism, and phenylpropanoid biosynthesis pathways (Table 2). Carbon partitioning is a critical process by which plants distribute the energy of photosynthesis and convert the assimilated carbon into sugar or its derivatives (Wang et al., 2013). Most plants store carbon as starch or cellulose (insoluble) with a lower concentration of sucrose (soluble), while sugarcane can store high concentrations of sucrose on its stems (Wang et al., 2013). Sucrose is cleaved into fructose and UDP-Glu, which is a nucleotide sugar precursor for most cell wall polysaccharides (Verbančič et al., 2018). Sugarcane maintains a dynamic balance of degradation of sucrose for respiration or its re-synthesis for storage. During this cycle, the carbon can be partitioned into other metabolites or fixed in polymers that can either be remobilized (such as starch in plastids) or added to structural biomass (such as cellulose, hemicelluloses, and lignin) (Wang et al., 2013). In this balance, the enzymes sucrose synthase (SuSy), sucrose phosphate synthase (SPS), sucrose phosphate phosphatase (SPP), and invertase play a central role in sucrose metabolism. At the same time, cellulose synthesis is catalyzed by enzyme complexes of cellulose synthase (CesA) (Stein & Granot, 2019). Cell wall biosynthesis can reduce sucrose accumulation since carbon fluxes directed to plant growth, and cell wall expansion may alter carbon partitioning into sucrose (Papini-Terzi et al., 2009). It is also possible that sucrose accumulation may trigger increased lignification (Papini-Terzi et al., 2009).

The STP assembly of the sugarcane genome has multiple copies of SuSy (11 copies), SPS (6 copies), SPP (2 copies), invertase (10 copies), and CesA (37 copies) annotated on its sequence. None of them is located in the top windows with higher heritability for the analyzed traits. However, window w194 (which explains 15% of h^2^_*G*_ for FB and 32.2% for TSH) contains a gene that encodes a copy of invertase (Sh04_g011120). Among the enzymes encoded by genes located in the top windows with higher heritability, phenylalanine ammonia-lyase (PAL) (Sh04_g015390, Sh04_g015400, and Sh04_g015410; w202 which explains 18% for PC) stands out as a critical enzyme involved in the phenylpropanoid pathway and biosynthesis of lignin (Zhang & Liu, 2015), which is also related to sucrose content (Papini-Terzi et al., 2009).

## 5. Conclusions

Throughout the analyses performed here, RHM has shown to be a useful approach to identify regions in the sugarcane genome related to agronomic traits. Even with the complexity of the sugarcane genome and its polyploidy impacting the identification of regions containing SNPs significantly associated with the phenotypes analyzed, the selection of windows that explained higher proportions of genomic heritability allows us to identify genomic regions containing a set of genes that are related to them. Among the selected windows, we identified a set of genes that encode enzymes that integrate metabolic pathways directly related to the traits analyzed. The selection of windows with higher heritability, therefore, represents an alternative for sugarcane breeders to reduce the complexity of the sugarcane genome since the selected windows span a region of 58.38Mb, which corresponds to 14,28% of the STP assembly of sugarcane genome. These windows correspond to promising genomic regions for the development of gene panels aiming the practice of marker-assisted selection of traits such as percentage of fiber in sugarcane bagasse (FB), apparent percentage of sucrose in sugarcane (PC), tonnes of pol per hectare (TPH) and tonnes of stalks per hectare (TSH). The findings obtained in this study will contribute to the progress of the genetic improvement of sugarcane.

## 6. Declarations

## Acknowledgments

The authors would like to thank the Núcleo de Análise de Biomoléculas (NuBioMol) and Sugarcane Breeding Program, Universidade Federal de Viçosa, MG, Brazil for providing the facilities necessary for the execution of the experiments.

## Funding

This work was funded by Coordenação de Aperfeiçoamento de Pessoal de Nível Superior (CAPES) - Finance Code 001, Conselho Nacional de Desenvolvimento Científico e

Tecnológico (CNPq), Fundação de Amparo à Pesquisa do Estado de Minas Gerais (FAPEMIG).

## Author’s contributions

PMPV, MM, GM, and LAP analyzed data and wrote the manuscript; MHPB and LAP planned and designed the research. PMAC conducted the experiments. MHPB and LAP coordinated the research. All authors reviewed and approved the manuscript.

## Supplementary Materials

### Supplementary Material 1. Tables

**Table S1. Adjusted phenotypic observations (genotypic values) of evaluated sugarcane clones.** This population consisted of 508 clones from 100 half-sib families, originated from crossings between elite sugarcane clones and commercial cultivars. Sugarcane clones were evaluated for the percentage of fiber in sugarcane bagasse (FB), apparent percentage of sucrose in sugarcane (PC), tonnes of pol per hectare (TPH), and tonnes of stalks per hectare (TSH).

**Table S2. Estimates of variance components and genetic parameters for percentage of fiber in sugarcane bagasse (FB), apparent percentage of sucrose in sugarcane (PC), tonnes of pol per hectare (TPH), and tonnes of stalks per hectare (TSH).** σ^2^_u_: genotypic variance. σ^2^_b_: variance between blocks. σ^2^_e_: residual variance. σ^2^_f_: individual phenotypic variance. H^2^: broad-sense heritability.

**Table S3. Windows with higher heritability in the sugarcane genome for analyzed traits**. h^2^_*r*_: window heritability. h^2^_*G*_: genomic heritability of the analyzed trait. h^2^_*r*_/ h^2^_*G*_: the proportion of genomic heritability explained by a window. Analyzed traits: percentage of fiber in sugarcane bagasse (FB), apparent percentage of sucrose in sugarcane (PC), tonnes of stalks per hectare (TSH), and tonnes of pol per hectare (TPH).

**Table S4. Functional annotation of genes included in top-5 windows with higher heritability in sugarcane genome for percentage of fiber in sugarcane bagasse (FB), apparent percentage of sucrose in sugarcane (PC), tonnes of pol per hectare (TPH) and tonnes of stalks per hectare (TSH).**

**Table S5. Gene Ontology (GO) terms lists summarized by REVIGO.** Non-redundant lists of GO terms assigned to the genes included in top-5 windows with higher heritability for percentage of fiber in sugarcane bagasse (FB), apparent percentage of sucrose in sugarcane (PC), tonnes of pol per hectare (TPH) and tonnes of stalks per hectare (TSH).

### Supplementary Material 2. Figures

**Figure S1. Schematic overview of the genotyping of sugarcane clones performed in this study.** The raw reads were processed, mapped to the sugarcane reference genome, and a variant calling for SNPs was performed. The software that was used for each step and their respective versions are indicated on the boxes. The command-lines used and the parameters which were considered are also shown.

**Figure S2. Significance analysis of RHM window-trait associations along the genome of analyzed sugarcane clones. (A).** Circular Manhattan plots (-log_10_ P) of RHM window-trait association for the percentage of fiber in sugarcane bagasse (FB), apparent percentage of sucrose in sugarcane (PC), tonnes of pol per hectare (TPH), and tonnes of stalks per hectare (TSH). **(B).** Quantile-quantile (QQ) plot of the data shown in the circular Manhattan plots.

