## Supplementary Material 2. Figures for "Regional genomic heritability mapping for agronomic traits in sugarcane"

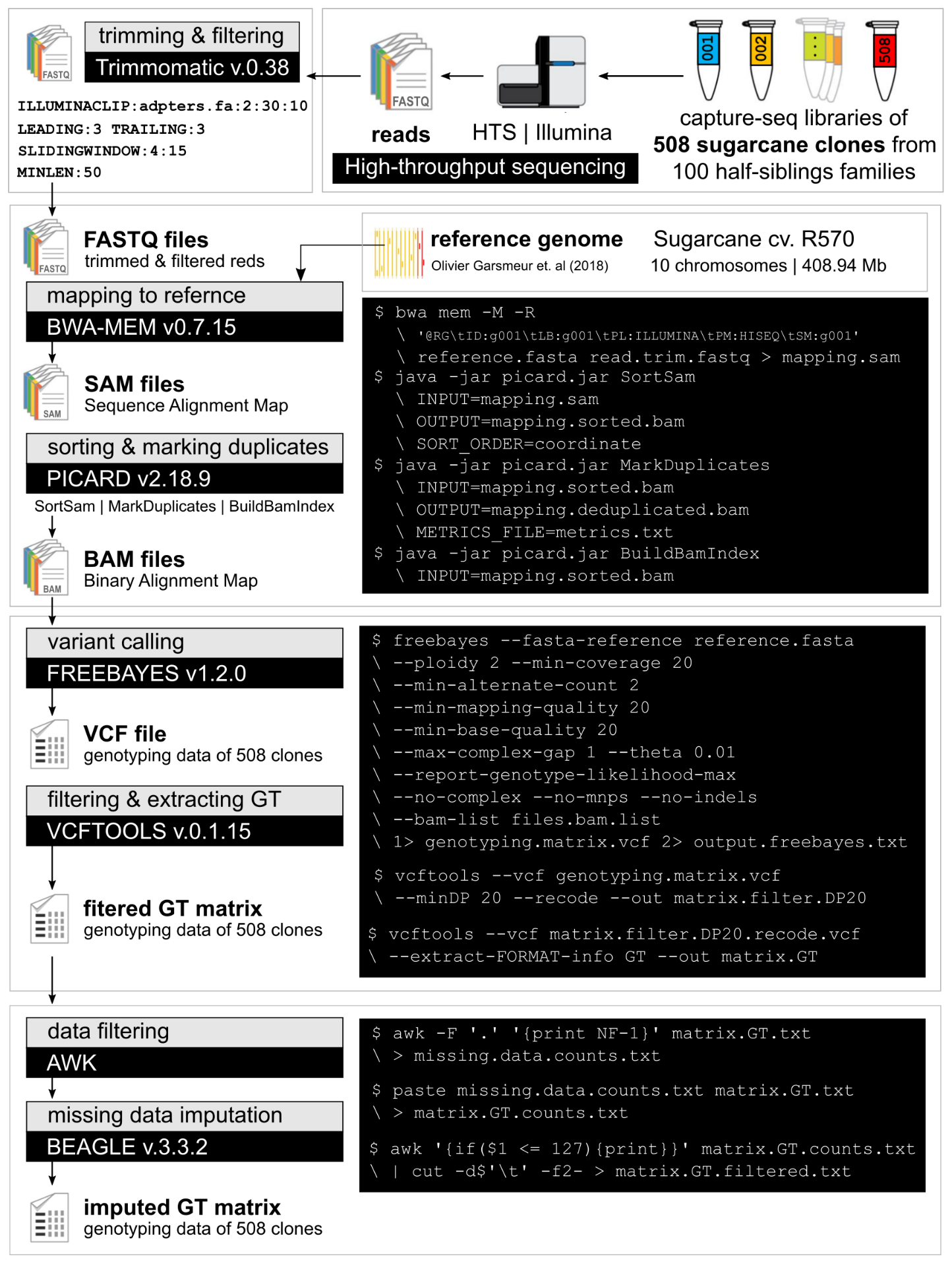

**Figure S1. Schematic overview of the genotyping of sugarcane clones performed in this study.** The raw reads were processed, mapped to the sugarcane reference genome, and a variant calling for SNPs was performed. The software which was used for each step and their respective versions are indicated on the boxes. The command-lines used and the parameters which were considered are also shown.

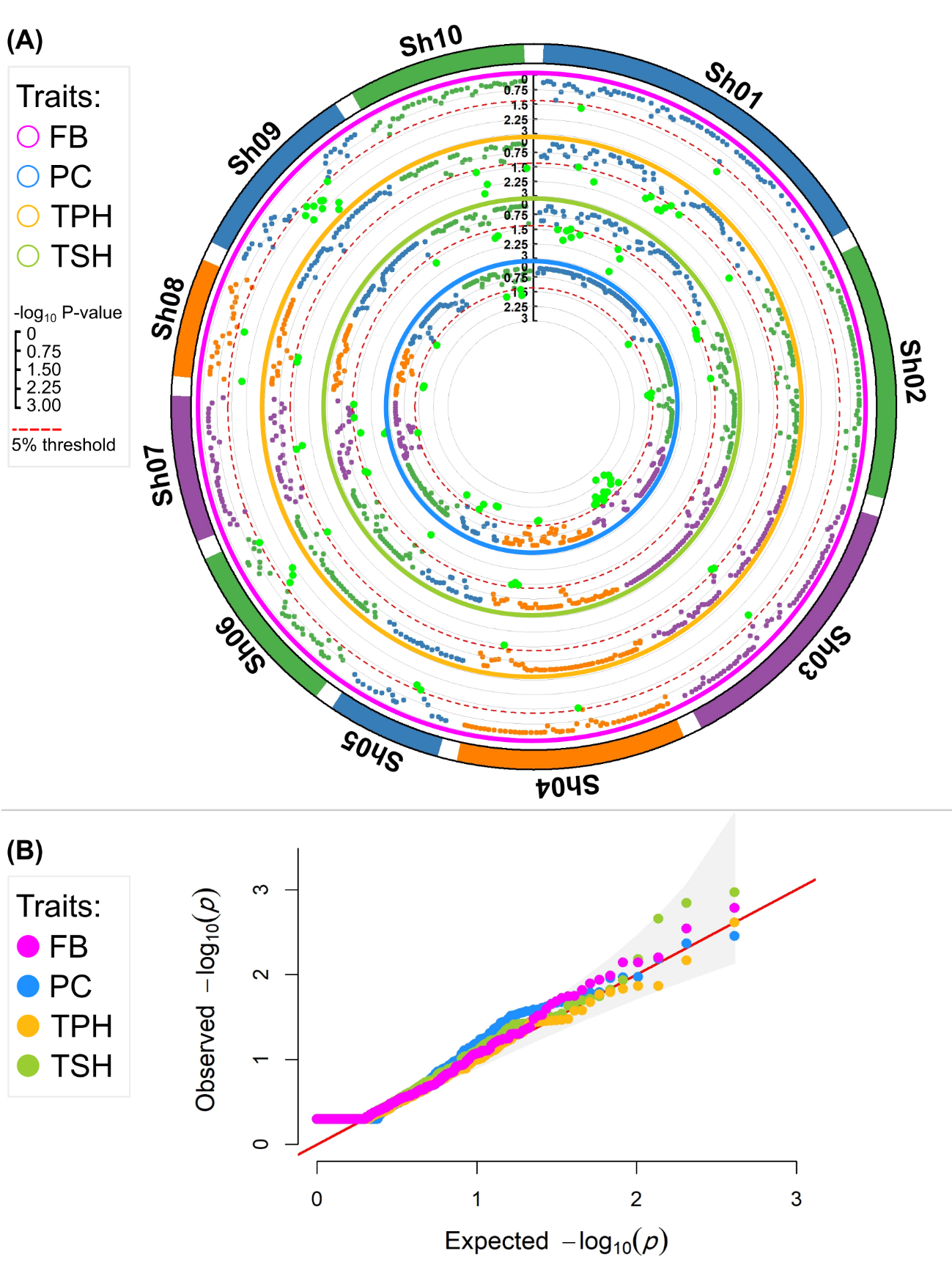

**Figure S2. Significance analysis of RHM window-trait associations along the genome of analyzed sugarcane clones. (A).** Circular Manhattan plots (-log_10_ P) of RHM window-trait association for the percentage of fiber in sugarcane bagasse (FB), apparent percentage of sucrose in sugarcane (PC), tonnes of pol per hectare (TPH), and tonnes of stalks per hectare (TSH). **(B).** Quantile-quantile (QQ) plot of the data shown in the circular Manhattan plots.
